# Adsorption Race Between Activated Carbon and Lactase for Intermolecular Interaction with Lactose

**DOI:** 10.1101/2022.09.29.510224

**Authors:** Emin Zümrütdal, Erdal Kuşvuran, Umut Kökbaş, Mine Çürük, Tuba Şimşek Mertoğlu, Büşra Dağ, Güray Kılınççeker

## Abstract

**Background:** Disturbing dyspeptic complaints may be seen in the use of milk and dairy products in people with lactose intolerance. Lactose-free milk and dairy products are produced for people with these complaints. The widespread use of activated carbon for dyspeptic complaints can also be used for adsorbing lactose.

**Methods:** For this purpose, the binding energy of lactase to lactose was studied in silico, lactose adsorption enthalpic changes of activated carbon were calculated by HPLC, plain and activated carbon yoghurt was produced and glucose+galactose and lactose levels were determined in yoghurts. The effects of these yogurts on serum glucose levels were compared in mice.

**Results:** In silico studies, the affinity of lactase with lactose was found to be -7.12 kcal/mol. It was determined by HPLC that activated carbon adsorbed lactose with an energy of -1,785 kcal/mol, and glucose+galactose levels and lactose ratios were lower in yogurt with added activated carbon. It was determined that there was no change in serum glucose levels in the 45th and 90th minutes following fasting in the mice fed with activated carbon yogurt compared to the mice fed plain yogurt.

**Conclusion:** Yogurt with activated carbon can be an alternative diet for individuals with lactose intolerance, by converting lactose to lactase in the presence of lactase and adsorbing lactose in the absence of lactose.

## Introduction

Carbon has been the building block of human beings since the existence of humanity, but it has also been an indispensable element for human survival. As science progresses, carbon continues to fascinate people with all its mysteries. While human life continues in the world, carbon has entered human life as coal, biochar, activated carbon and graphene(1).

While these different forms of carbon find a place for themselves in various industries, the adsorbent property of activated carbon has been used for water and food consumption(2,3). Although activated carbon is used in emergency services in cases of poisoning, activated carbon and biochar have been observed and used in many clinical conditions such as the excretion of toxins from the body and irritable bowel syndrome(4,5,6). One of these studies is yogurt with activated carbon, which was created with the aim of benefiting from the lactose adsorption effect of activated carbon(7,8).

While the consumption of yogurt is common in the Middle East countries, its use has increased rapidly all over the world for the last half century. Yogurt gains nutritional properties with the protein, fat, sugar, minerals and microorganisms it contains(9). Protein, fat, carbohydrates and minerals in yogurt are very important for nutrition in newborns and childhood. Humans take carbohydrates from different foods in their adult life. Therefore, protein, fat, minerals, Lactobacillus Bulgaricus and Streptococcus Thermofilus in yogurt for adults come to the fore. In order for lactose, which is a carbohydrate from milk, to be digested in yogurt, lactose must be broken down into glucose and galactose by the lactase enzyme. Glucose and galactose are absorbed from the small intestine and enter the systemic circulation (10).

However, yoghurt consumption may bring along some dyspeptic complaints(11). The biggest factor causing dyspeptic complaints in yoghurt consumption is lactose, which is usually found in yoghurt in lactose intolerant individuals (12).

Lactose intolerance, which is very common, is associated with many reasons, especially the lifestyle of people and the geographical regions they live in(13). While one third of adults maintain their ability to digest lactose, 2/3 of them have digestive complaints varying in severity depending on the person.

Lactose-free yogurt has been produced in order to continue the use of yogurt, which is very beneficial for people with lactose intolerance. With this method, milk is treated with lactase enzyme. Lactase in milk breaks down lactose into glucose and galactose (14). Since there is no lactose in milk, milk and yogurt obtained from this milk are used as lactose-free products(15).

In this multidisciplinary study, the ability of activated carbon to adsorb lactose was evaluated by in silico study. In order to support these findings, lactose adsorbance findings of activated carbon were studied by HPLC. In order to support in silico and HPLC studies, the effect of activated carbon on lactose digestion was investigated by in vivo experiments in mice.

## Methods

### Ligand and Receptor retrieval

In this molecular docking study, beta-galactosidase enzyme (E. coli (LACZ) beta-galactosidase (E537Q) in complex with lactose) used as a target receptor was downloaded from the RCSB PDB database (https://www.rcsb.org/) with the code 1JYN. Prior to docking calculations, water molecules, non-interacting ions and co-crystallized ligand (lactose) were removed from the protein structure. Lactose, which was docked as a ligand against the beta-galactosidase enzyme in the study, was extracted computationally from the 1JYN complex and further utilized accordingly. Thus, the binding strength of the lactose molecule, which occupies the active site of the enzyme as a ligand (substrate) in the crystallized binary complex, was calculated by molecular docking with the beta-galactosidase enzyme.

### Molecular docking between beta-galactosidase and lactose

Prior to docking simulations with AutoDock Vina, target (beta-galactosidase) and ligand (lactose) structures were prepared using AutoDockTools 1.5.6 and saved in pdbqt format (16,17). In molecular docking simulations with E. coli beta-galactosidase and lactose, polar hydrogen atoms in the receptor and ligand molecule were retained, whereas, non-polar hydrogens were merged. Kollmann charges were assigned to the receptor and Gasteiger charges were assigned to the lactose. In addition, torsional degrees of freedom of the ligand’s (lactose) rotatable bonds were allowed in this docking protocol. A grid box size of 40×40×40 Å points (x = -16.69; y = -31.08; z = -26.55) and a 0.375 Å grid spacing was defined as the search space for putative interactions between the enzyme and ligand. This grid box size was adjusted in a way that the lactose molecule could easily interact with the side chains of active site residues of the beta-galactosidase. After 20 independent docking runs of lactose against the beta-galactosidase, all potential binding modes of the ligand were clustered by AutoDock Vina on the basis of geometrical similarity and were ranked based on the binding free energy (ΔG°; kcal/mol) of each of the ligand conformations which showed the lowest (best) affinity against the target receptor. The top-ranked docking conformations (modes) of the lactose molecule calculated by AutoDock Vina were rendered and analyzed using Discovery Studio Visualizer v16 (18,19).

### Production of Plain and Activated Carbon Yogurt

Plain and activated carbon added yoghurt trials were carried out according to the methods described previously(8,20). Milk (pasteurized, semi-skimmed, homogenized cow’s milk) milked on the same day was used in the study (Milk was obtained from Çukurova University Faculty of Agriculture Research and Application Farm livestock branch). A total of 6 L of milk was put into 2 separate boilers with 3 liters each. An additional 3 g of activated carbon (Medical Charcoal, CAS No:7440-44-0, Production date:01/2021, BATCH NO:2102H0060, GALENİK) was added into a boiler and mixed for 4 minutes.

Milk in both groups was pasteurized at 90 °C. Plain milk was left to cool to ferment. After the milk with activated carbon was taken from the stove, it was filtered through a sterile milk filtration filter a total of 3 times, once at 15 and 30 minutes. A starter (Danisco, Yo Mix 505 LYO 200 DCU-France) was used in both groups at 47 °C at a rate of 3% (0.078 g per 3 L). After the starter was added, the milk was mixed gently for 4 minutes and then left for 4 minutes. Then the milk was placed in standard yogurt cups (200mL standard yogurt cups). After this process, the milk is placed in ovens (47±2 °C) to be incubated.

The products that had the consistency of yogurt were checked and removed from the oven after 4 hours and 10 minutes. The mouths of the containers were closed and removed to +4 °C cold storage without waiting. All operations were carried out in the Çukurova University Faculty of Agriculture Research and Application Farm Food branch dairy business.

### Measurement of Lactose-Activated Carbon Interaction via HPLC

The efficiency of adsorbing of lactose on acitvated carbon was determined with the help of Langmuir adsorption isotherm(21,22). First, 0.500 g of activated carbon was placed in to the 15 mL polypropylene Falcon tubes with screw caps. Six different concentrations of lactose solutions were prepared and a portion of the solution was separated to two portion. One of them was used to determine the initial concentration. Another 10 mL portion of lactose solutions was added to the Falcon tubes activated carbon contained. Seperatly it was shaken overnight in a shaking water bath at temperatures of 15 °C, 25 °C and 35 °C. Before it was centrifuged at 3000 rpm for 10 minutes (Nüve Brand-Turkey).

Then, 20 μL of the supernatant activated carbon free of supernatant from activated carbon was injected into the HPLC device (Shimadzu brand 10A RID, Japan) equpmented with reflactive index detector. Equilibrium concentrations of lactose solution in adsortion-desorbtion experiments were determined by HPLC analysis with 70% Acetonitrile-Water mixture mobile phase and flow rate of 0.8 mL/min. On the other hand a portion 10 mL was analyzed by HPLC also. When the obtained results were evaluated, it was determined that there was a linear relationship with the Langmuir adsorption equation (Equation 1) (Figure 1).

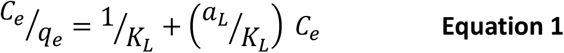

Lactose, glucose and galactose values in yogurt samples were evaluated as following procedure. Yogurt samples with and without activated carbon were homogenized for 5 minutes with vortex. It was centrifuged at 3000 rpm for 10 minutes in the centrifuge device. For each sample, 20 μL of the clear supernatant was taken and injected into HPLC.

**Figure 1.**
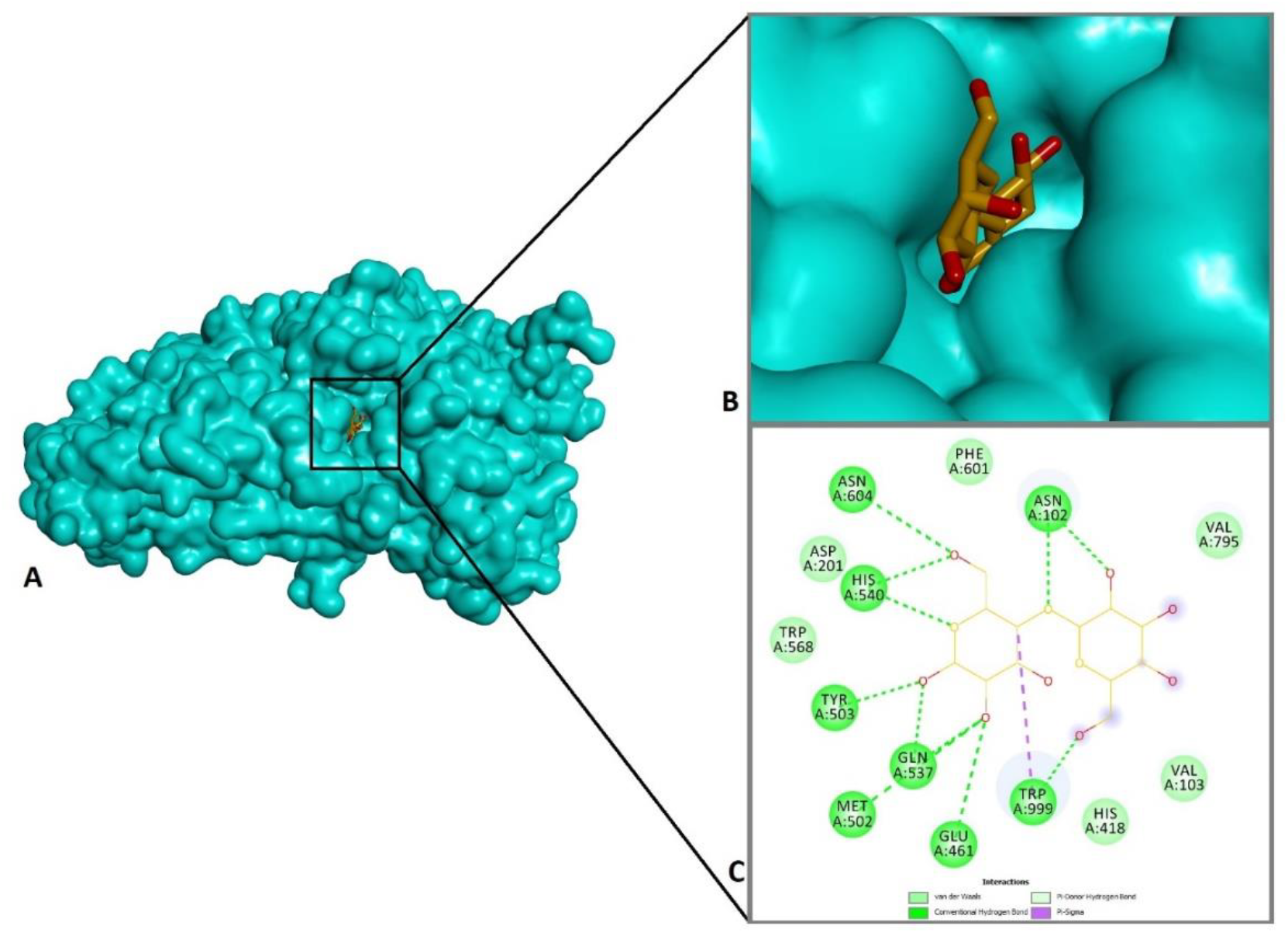
(A) 3-dimensional general view of the docking complex calculated between the lactose molecule and the beta-galactosidase (=lactase) enzyme. Lactase enzyme is shown in the molecular surface (turquoise) and the ligand (yellow) is shown in the stick mode. (B) The zoom in image of the predicted docking conformation and orientation of the lactose molecule within the catalytic pocket of the enzyme. Note that this pocket which forms the active site of the enzyme is relatively narrow in volume. (C) The 2D image of the interactions of lactose with the amino acid residues in the active site of the lactase enzyme. Green dashed lines represent hydrogen bonds and purple dashed lines represent hydrophobic contacts. Peripheral amino acids (Val, Phe, Asp, Trp and His) that do not participate in bond formation, however, constitute the van der Waals surface of protein-ligand interaction (see the 2D image, C).

### Ethics committee decision

The study was taken with the decision of the Ethics Committee of Adana Veterinary Control Institute, Animal Experiments Local Ethics Committee (Adana/Turkey). Date: 05.09.2022, Decision no: 2022-7.

### Measurement of serum glucose ratios in animals

After 12 hours of fasting, at 08:00, serum glucose values of all groups were determined by glucometer from the tail blood after anesthesia. Then, 1 mL of drinking water was given to the 1st group, 1 mL of plain yogurt to the 2nd group, and 1 mL of activated carbon yogurt to the 3rd group by gavage. Afterwards, changes in serum glucose values were monitored at 08.45 and 09.30 hours (23). (Procedure details are in the supplement file.)

### Statistical evaluation

Statistical analyzes were performed with the SPSS software (SPSS, version 17.0, IBM Corporation, Armonk, NY, USA). The paired samples t test was used to analyze the numeric data of the serum glucose level of mice and glucose+galactose and lactose values in plain yogurt and yogurt with activated carbon. The value of p<0.05 was considered to be statistically significant.

## Results

### Molecular interactions of lactose with the beta-galactosidase enzyme

In this molecular docking study, the molecular affinity of the lactose molecule against the beta-galactosidase enzyme was determined (Table 1), and the non-bonded interactions in this binary complex between the enzyme active site and the lactose (ligand) were shown (Figure 1C). Considering the lactose molecule is the native substrate of beta-galactosidase, our docking study is a confirmation in which the binding strength and the interactions of this ligand against the beta-galactosidase was re-calculated.

**Table 1.** Beta-galactosidase non-bonded interaction results for lactose and the predicted docking binding free energy (ΔG°) of the formed complex.

| Compound | Receptor | $\Delta G_{\text{best}}$<br>(kcal/mol) | Classical<br>H-bond | Hydrophobic<br>(pi-sigma) |
| --- | --- | --- | --- | --- |
| Lactose | 1JYN | -7.12 | Asn102, Glu461,<br>Met502, Tyr503,<br>Gln537, His540,<br>Asn604, Trp999 | Trp999 |
| Total number<br>of non-bonded<br>interactions |  |  | 24 | 2 |

1JYN (receptor): *E. coli* (LACZ) beta-galactosidase (E537Q) in complex with lactose
$\Delta G_{\text{best}}$ : most favorable binding free energy (kcal/mol).

Lactose formed multiple hydrogen bonds with residues Asn102, Glu461, Met502, Tyr503, Gln537, His540, Asn604, and Trp999 at the active site of the beta-galactosidase, as well as additional hydrophobic pi-sigma contacts with the residue Trp999 (Table 1, Figure 1C). In addition, the binding free energy of the formed complex was found to be highly favorable (ΔG°=-7.12 kcal/mol, Table X). Taken together, it is evident that the molecular complex formed between lactose and beta-galactosidase is stabilized by a large number of H-bonds (24 in total, Table 1 and Figure 1C), and this calculated huge number of interactions proves the success of the docking protocol: Docking conformational search algorithm can produce poses close to the chemically most favorable conformation of the ligand.

### Lactose-Activated Carbon Interaction

The lactose solutions prepared to use in adsorbtion-desorbtion studies were observe to seperated to different concentration values such as theorical and experimental after HPLC analysis. The experimental concentration values were accepted as initial concentration values. Theoritical, initial and equilibrium concentrations values can be seen Table 2. While the amount of adsorbed lactose was increasing with increasing of the initial concentration, at every 3 temperature, the lactose percentage of lactose concentration adsorbed on activated carbon were decreasing with increasing all the initial concentration. In the adsorption study at 15 °C, the amount of lactose adsorbed at the highest concentration was 8.3%, while it was 37.2% at the lowest concentrationWhen the adsorbed amounts were compared,2405 mgL^-1^lactose was adsorbed at the highest concentration, while 1826 mgL^-1^ lactose was adsorbed at the lowest concentration.

**Table 2.**
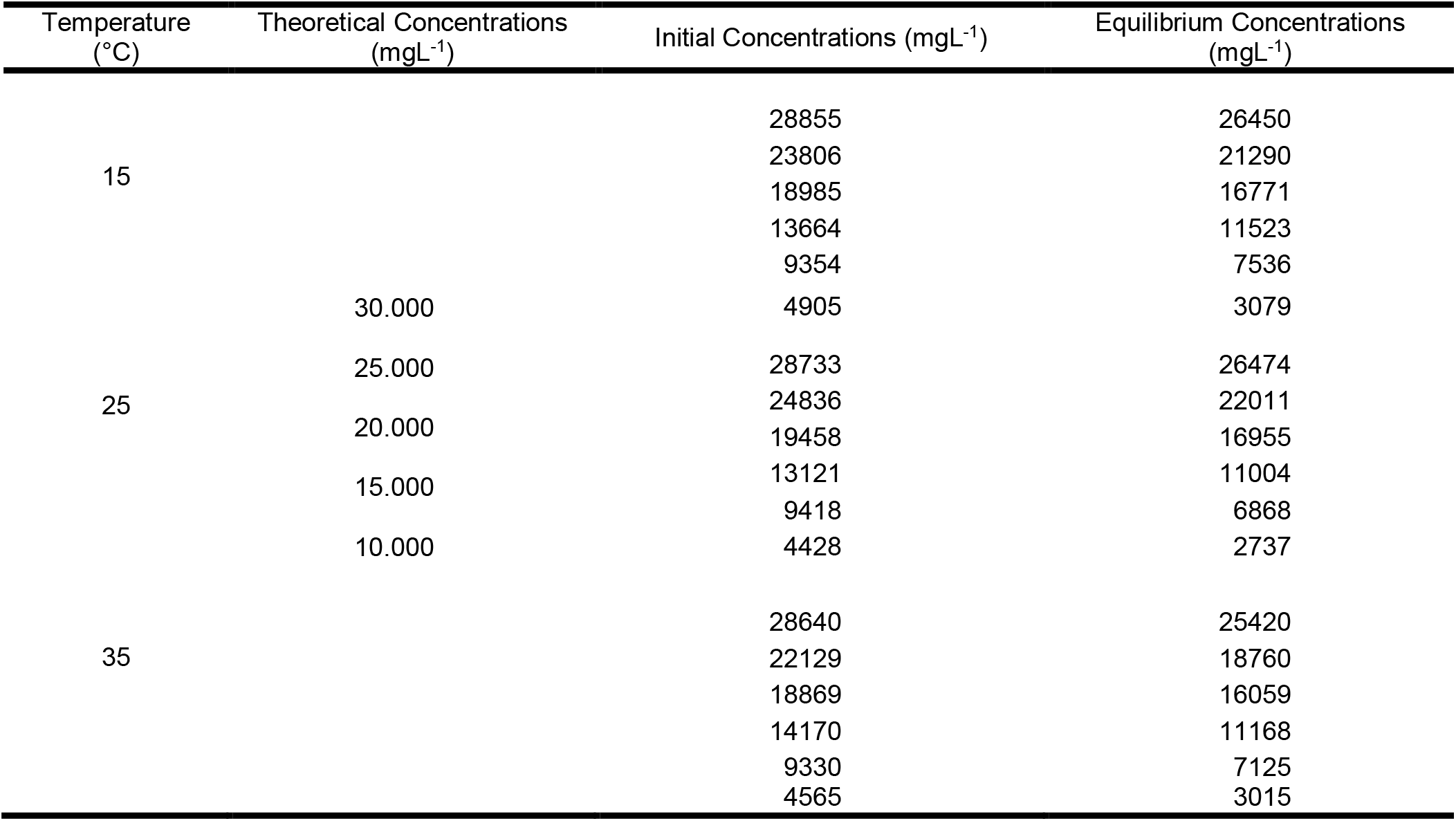
Lactose theoretical concentrations, lactose initial concentrations and lactose equilibrium concentrations for Lactose-Activated Carbon Adsorption

The regression coefficiency of the lines obtained at 15 °C, 25 °C and 35°C temperatures were found to be 0.994, 0.995 and 0.988, respectively (Figure 2). Maximum adsorption values qmax and equilibrium constant values are given in Table 3.

**Figure 2.**
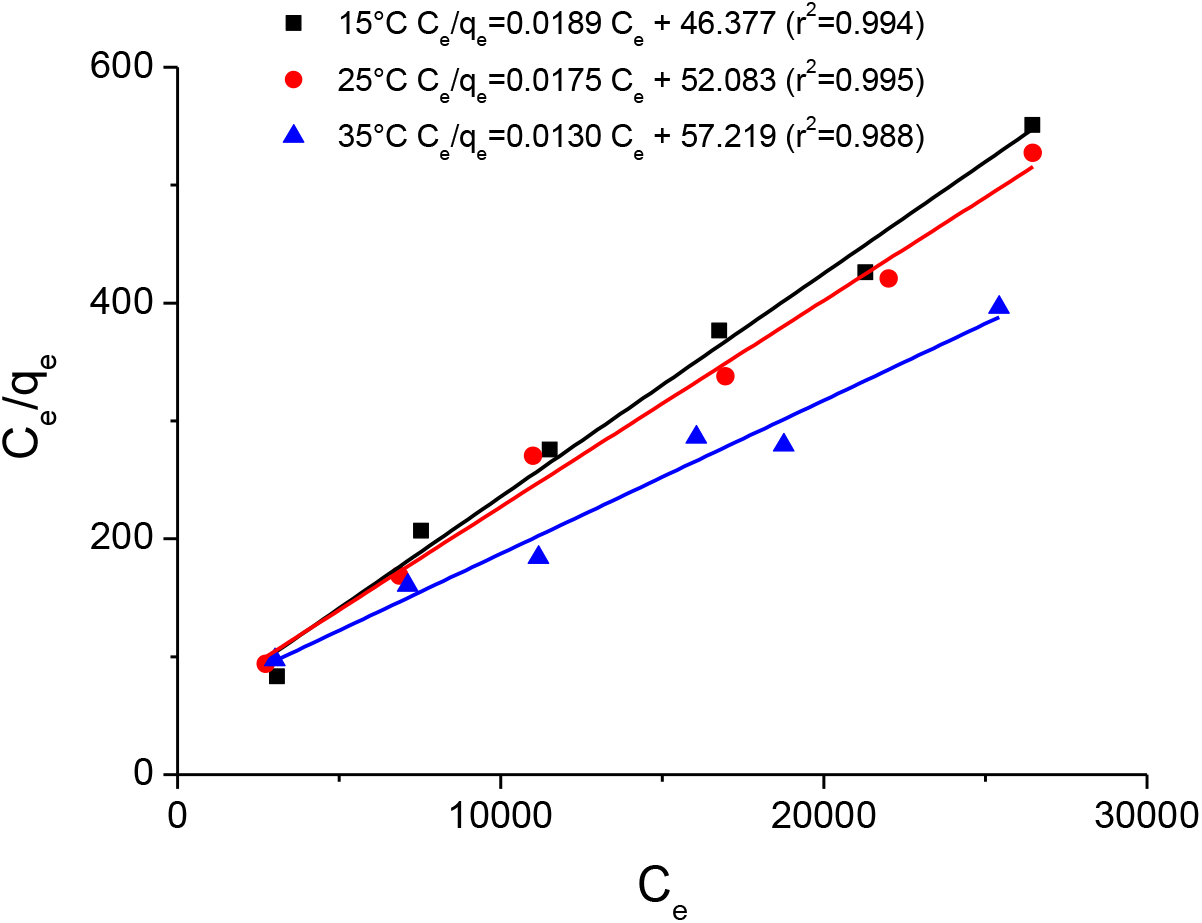
Lactose-Activated Carbon interaction at 15°C, 25°C and 35°C Langmuir adsorption isotherms. Ce: equilibrium concentration (mg/L), qe: amount of Latose adsorbed on Activated Carbon (mg/g).

**Table 3.** Lactose-Activated Carbon adsorption Langmiur adsorption equation parameters

| Temperature,<br>°C | $a_L/K_L$ | $K_L$ | $q_{max}$ (mg/g) |
| --- | --- | --- | --- |
| 15 | 0.0189 | 0.0214 | 52.9 |
| 25 | 0.0175 | 0.0192 | 57.1 |
| 35 | 0.0130 | 0.0175 | 76.9 |

The relationship between the adsorption-desorption equilibrium constant and the temperature is as below.

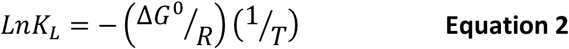

According to Equation 2 when the equilibrium constants are plotted versus 1/T values, the slope of the line will be equal to Δ*G*^0^*/R* (Figure 3).

**Figure 3.**
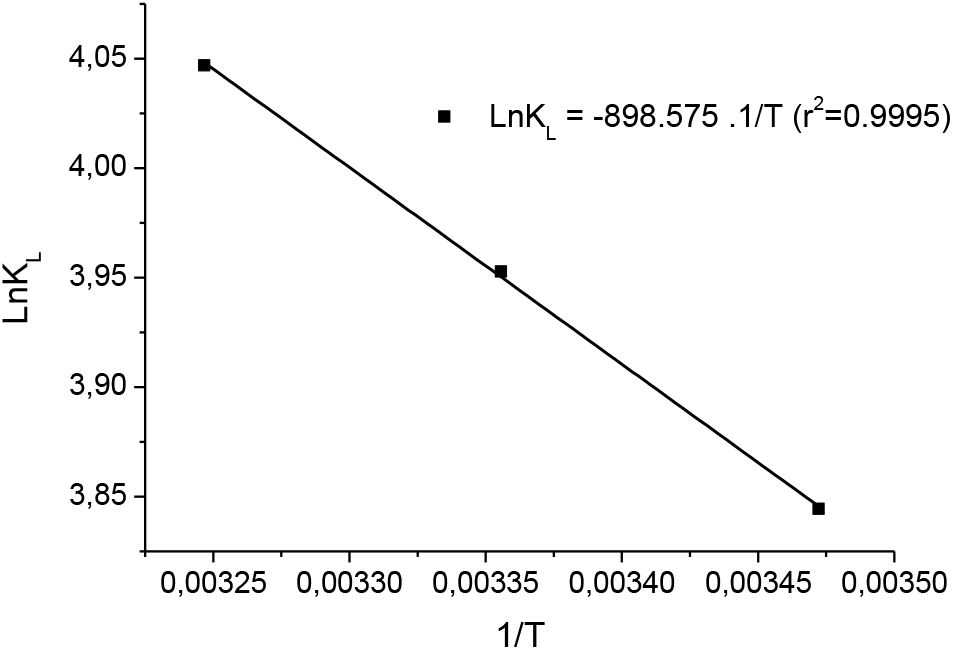
Changes versus 1/T values

According to Equation 2, the free enthalpy change was found to be – 1.785 kcal/mol.

### Determination of glucose, galactose and lactose amounts in plain yoghurt and activated carbon added yoghurt

Glucose, galactose and lactose values were determined by HPLC in plain yoghurt and activated carbon added yoghurt. In the evaluation, glucose+galactose(p<0.05) and lactose(p<0.01) values were found to be statistically higher in plain yogurt compared to yogurt with activated carbon (Figure 4). (Glucose+galactose and lactose values measured by HPLC for plain yoghurt and activated carbon added yoghurt are given in the attached file.)

**Figure 4.**
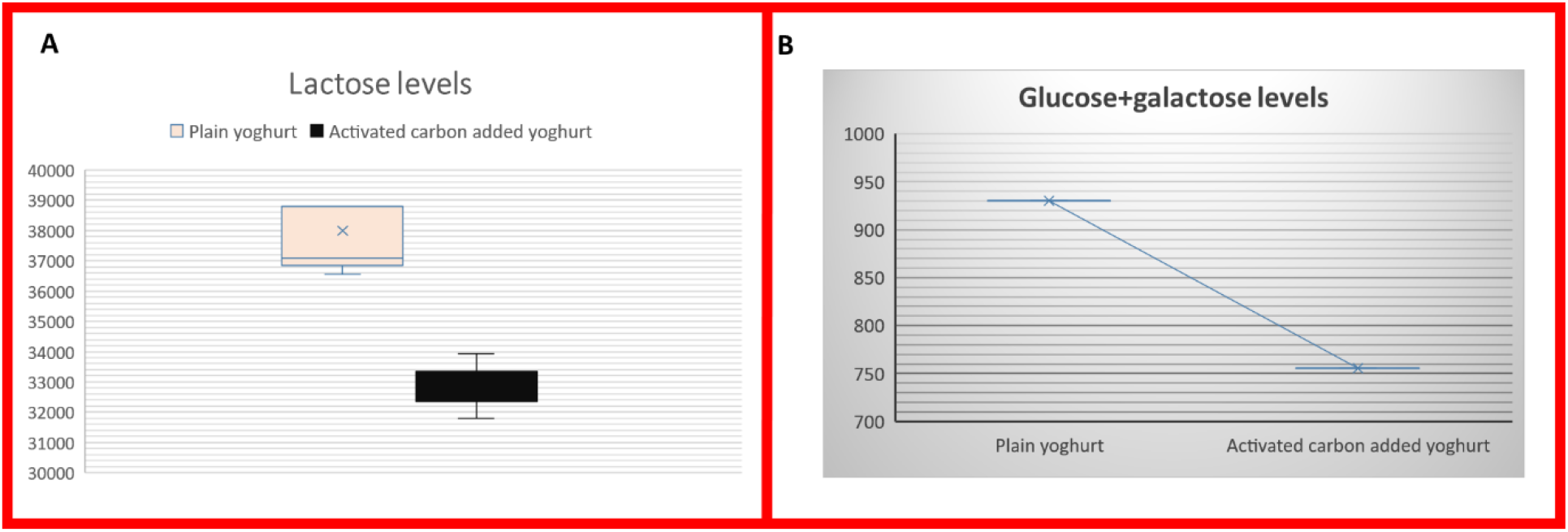
Glucose, galactose and lactose amounts(mg/L) in plain yogurt and activated carbon added yogurt according to HPLC measurements

### Serum glucose levels

Serum glucose values obtained in the groups did not differ in all groups after 12 hours of fasting. No statistical change was observed in serum glucose values obtained in the control group at the 45th and 90th minutes. However, the increase in serum glucose values in the subjects fed with plain and activated carbon added yoghurt was statistically significant compared to the control group. No statistically significant difference was observed between serum glucose values at 45th and 90th minutes in the groups fed plain yogurt and yogurts with activated carbon added (Figure 5). given.)

**Figure 5.**
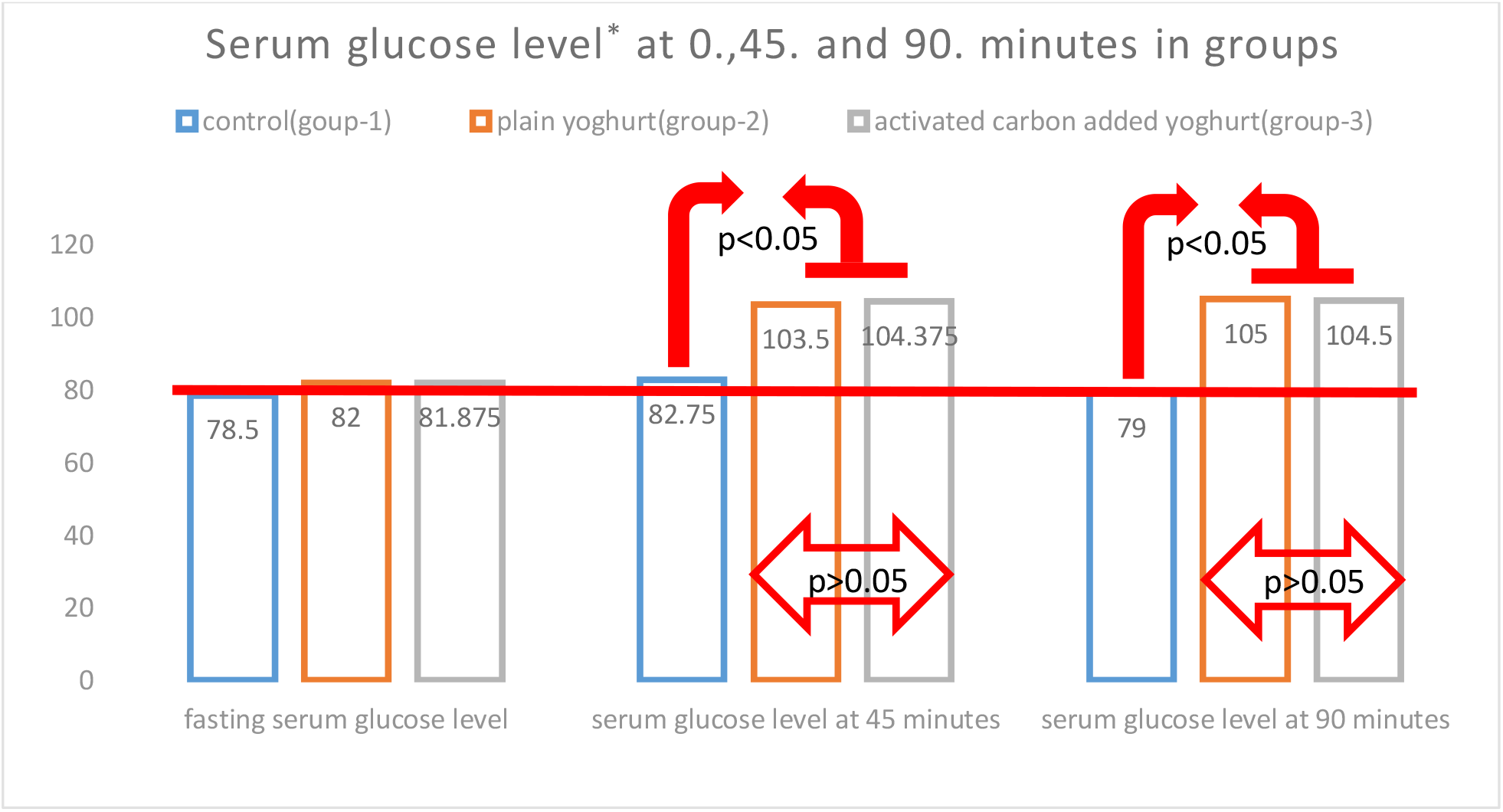
Serum glucose levels (*mg/dL).

## Discussion

In this study, the interaction forces between lactase and activated carbon to bind lactose and its clinical reflection were evaluated and presented for the first time in the literature, according to our knowledge.

In the previous study, the interaction energy of -4.01 kcal/mol was determined between lactose and activated carbon(4). In this study, the interaction energy between lactose and lactase was evaluated and this value was found to be -7.12 kcal/mol. This energy difference suggests that lactose molecule and lactase enzyme can interact more easily than activated carbon.

The consumption of yoghurt in adults gains value especially because of the protein, fat, minerals and beneficial microorganisms it contains. In lactose intolerance, lactase enzyme deficiency causes lactose to be perceived as a foreign body in the intestines. This situation leads to troublesome dyspeptic complaints with the accumulation of water in the intestinal lumen in the small intestines, then the accumulation of methane, carbon dioxide and hydrogen gases in the large intestines due to the use of lactose by the microorganisms. For these reasons, lactose in the lactose-free yogurt developed is previously broken down by the lactase enzyme. However, in these yogurts, glucose and galactose values, which can be absorbed very easily into the blood, increase in yoghurt with lactose breakdown compared to normal yogurt.

In the study, activated carbon was used in lactose intolerance to transport lactose from the intestines without being recognized as a foreign body.

In studies with HPLC, the free enthalpy exchange energy after the interaction of activated carbon with lactose was found to be -1.785 kcal/mol according to equation 2. This result had the same features as the previous docking study result(4). However, this result supported the previous docking work but contained different value. It was thought that this difference might be due to the fact that the docking operation was created in an environment without water. The increase of chemical potential of lactose solution depend on increase of concentration of lactose will force that more lactose will be adsorbed on activated carbon. In our study, we used 1 gram of activated carbon for 1L of milk. These values may differ with larger studies on the concentration of activated carbon in milk.

In the lactose measurements made by HPLC in yoghurts, the fact that lactose ratios are lower in yoghurts containing activated carbon than plain yoghurt makes us think that the lactose adsorbing capacity of activated carbon is correct (Figure 4).

In the same study, the low glucose and galactose levels in yogurts with added activated carbon were consistent with the difference in lactose amounts.

Lactose is a disaccharide. It is broken down into glucose and galactose monosaccharides by the enzyme lactase in the intestines. Glucose and galactose are rapidly absorbed from the intestines into the blood. In the study on in vivo serum glucose levels, there was no difference between serum glucose levels at the 0th minute after 12 hours of fasting in all groups that were given only drinking water, plain yogurt, and yogurt with added activated carbon (Figure 5).

There was no difference between the serum glucose levels at the 45th and 90th minutes compared to the 0th minute in the group given only drinking water. Since no energy source was given to the body, this was accepted as the expected value (Figure 5).

In the group given plain yogurt and yogurt with activated carbon, an increase in serum glucose levels was found at the 0th minute, 45th and 90th minutes. It was thought that this increase came from glucose, which is the breakdown product of lactose in the yogurt given to the body. This finding also shows the presence of lactase enzyme in the intestines of the experimental animals used.

The lack of difference in serum glucose levels between plain yogurt and yogurt with added activated carbon suggests that lactase interacts by separating lactose from activated carbon when there is lactase enzyme in the body. The higher affinity of the lactase enzyme, which was shown in the in silico study, to the catalytic pocket of lactose compared to activated carbon, correlates with serum glucose values. The interaction of the molecular complex formed between lactose and beta-galactosidase with multiple H-bonds (24 in total, Table 1 and Figure 1C) strengthens the affinity.

In previous studies, it has been shown that the addition of activated carbon does not affect the protein, fat, lactic acid, pH, Lactobacillus bulgaricus and Streptococcus thermophilus values that give yogurt its properties. In addition, it was determined that lactic acid values were lower and pH was higher in activated carbon yoghurt, although it remained within the spectrum range suitable for yoghurt properties. It was shown that activated carbon did not affect the calcium level, no difference was observed in the texture structure in the aroma test, however, it was better in general likability than plain yogurt.

## Conclusion

These results show that lactose is adsorbed by activated carbon, but when lactase is present, lactose interacts with lactase.

With these results, yoghurt with activated carbon may be an alternative to lactose-free yogurt, which does not contain lactose but contains high amounts of glucose and galactose, for people with lactose intolerance. Considering the effect of activated carbon on the excretion of toxins in the body, this alternative can be applied more effectively.

## Supporting information

It will not be added to the manuscript page of the article. This file contains descriptive information.

It will not be added to the manuscript page of the article. This file contains descriptive information.

## Acknowledgement

We would like to thank Gülden Sevgen in the anesthesia applications in the animal study and Erman Salih Istifli in the docking studies for their assistance.

