## Supplementary material for "Adsorption Race Between Activated Carbon and Lactase for Intermolecular Interaction with Lactose": It will not be added to the manuscript page of the article. This file contains descriptive information.

**Glucose+galactose level in yoghurt**

| | Plain Yoghurt(mg/L) | Yoghurt with activated carbon(mg/L) |
|---|---|---|
| | 887 | 703 |
| | 868 | 810 |
| | 863 | 741 |
| | 830 | 720 |
| | 881 | 731 |
| | 1251 | 827 |
| **Mean±SD** | **930±158,51** | **755,33±50,81** |

**Lactose level in yoghurt**

| | Plain Yoghurt(mg/L) | Yoghurt with activated carbon(mg/L) |
|---|---|---|
| | 37155 | 33138 |
| | 36994 | 31801 |
| | 37439 | 32570 |
| | 36545 | 32814 |
| | 36951 | 32554 |
| | 42877 | 33931 |
| **Mean±SD** | **37993,50±2410,11** | **32801,33±707,81** |
