## Supplementary material for "Adsorption Race Between Activated Carbon and Lactase for Intermolecular Interaction with Lactose": It will not be added to the manuscript page of the article. This file contains descriptive information.

**Animal experiments**

The experiment was performed on 24 Swiss Albino, 8 weeks old (25-30 g) male mice. Three groups (n=8) were formed in the experiment. Before the study, animals were fed 6.5g of standard mouse chow per mouse for 5 days and drinking water was given continuously.

The animals were kept at the same room temperature (25°C) for 12 hours in light and 12 hours in darkness throughout the experiment. Mice were fasted for 12 hours the day before the experiment. During this period, the cages of the animals were renewed so that no feed residues were left in the cage.

Each animal's ear was numbered by painting. Changes in serum glucose in each animal were noted separately. 15 minutes after the animals were anesthetized, the tail was cut 0.5 cm and serum glucose values were determined with a glucometer (GlucoDr/allmedicus, Germany). Serum glucose values were determined in all groups at 08:00 after 12 hours of fasting. The animals were anesthetized by administering intramuscular ketamine 150 mg/kg through the gluteal region. Then, 1 mL of drinking water was given to the 1st group, 1 mL of plain yogurt to the 2nd group, and 1 mL of activated carbon yogurt to the 3rd group by gavage. Afterwards, changes in blood glucose values were monitored at 08.45 and 09.30 hours.
